# Soil stockpile age does not impact vegetation establishment in a cold, arid natural gas field

**DOI:** 10.1101/2023.12.01.569687

**Authors:** Michael F. Curran, Joshua R. Sorenson, Timothy J. Robinson

## Abstract

Land reclamation is critical to ensure surface disturbance associated with natural gas development is not permanent. Soil management is critical to reclamation success, especially in environments which are challenged by aridity. Typically, natural gas well pad construction involves stripping topsoil to allow for equipment to be on level ground and placing it into a stockpile. After well pad construction is complete, soil is respread and seeded across a large portion of the initial disturbance to initiate interim reclamation. With the advent of new extraction technologies (e.g., directional drilling), it is not always known how many individual wells will be placed on a given well pad until results from exploratory tests are examined. Previous research has shown soil disturbance during natural gas well pad construction and subsequent reclamation in cold, arid environments is highest at the stripping and respreading phases, with minimal soil activity occurring during the stockpile phase. Other research has shown that additional soil disturbances after reclamation is initiated may exacerbate soil damage, limiting revegetation potential. However, no studies have been conducted to determine if soil stockpile age impacts vegetation emergence. Here, we examine soil stockpiles which are 1-7 years old in the Jonah Infill natural gas field for vegetation emergence and vegetation cover using an image analysis software called SamplePoint. In a ten-week greenhouse experiment, we found vegetation cover across stockpile age-classes increased uniformly during the study period but that there was no significant difference in rate of vegetation cover increase or percent vegetation cover over time. These findings suggest it may be better to keep soil stockpiled in cold, arid natural gas fields when it is uncertain if additional construction activities will be required on a well pad location rather than respreading soil with a chance that redisturbance is necessary.

## Introduction

Land reclamation and habitat restoration efforts are critical to ensure land surface disturbance associated with natural resource extraction does not have permanent negative impacts in Wyoming (Stahl & Curran 2017). Given that there is a high risk of failure associated with reclamation in arid environments, studies to understand and improve management practices are essential (Shackelford et al. 2021). Reclamation success in Wyoming oil and natural gas fields is often measured by above ground vegetation characteristics such as species richness or percent vegetation cover as a proxy for erosion control (Curran et al. 2013; Curran & Stahl 2015). However, soils are critical to the success or failures of reclamation outcomes and soil management is paramount to vegetation outcomes of reclamation efforts (e.g., Heneghan et al. 2008).

In Wyoming’s oil and gas fields, suitable topsoil depth is identified prior to well pad construction and topsoil is then stripped and stockpiled adjacent to disturbed areas and subsequently respread after exploration activities are complete (DOI 2007). While this disturbance may be significant, soil management related to oil and gas well pad reclamation typically does not include challenges associated with mining such as mixing with overburden or severe recontouring during the respreading phase. Nevertheless, understanding how soil management and handling impacts vegetation recovery in oil and gas fields is critical to improving reclamation success but is not well studied. In an annotated bibliography containing peer-reviewed literature, technical reports, theses and dissertations, and other grey matter which may be relevant to oil and gas reclamation in the western US, <10% of 300 documents examined soil stockpiles (Mann et al. *In Press*). Within those few studies, the majority examined soil characteristics or processes, and most were related to mining rather than oil and gas with little emphasis given to how vegetation responds to various soil stockpile management techniques (Mann et al. *In Press*).

In the Jonah Infill natural gas field of Sublette County, WY, USA (hereafter; Jonah Field), extraction had been primarily in the form of single vertical well pads until the last decade. In this scenario, well pads were generally built, drilled, and brought into production within a single season, allowing reclamation to occur within a year of breaking ground. With advancements in drilling technology (i.e., directional drilling), operators in the Jonah Field have transitioned to larger multi-well pads which may take more than one year to develop, therefore increasing the time necessary for soils to be stockpiled. Previous research in the area suggests major alterations in soil occur during the soil stripping phase or during the respreading phase with negligible changes occurring while soil remains in the stockpile (Mason et al. 2011). Since soils in stockpiles may lose biota dependent on plant carbon inputs if stored for long durations (e.g., Valliere et al. 2022), stockpiles in the Jonah Field are seeded to prevent erosion and maintain soil health within stockpiles during storage.

As soil handling is one of the most expensive costs of reclamation associated with natural gas development (e.g., Chenoweth et al. 2012), understanding how vegetation can establish from stockpiles of various ages is critical to operators. In this study, we examined soil stockpiles ranging from 1-7 years old. We utilized a seed mix common to the Jonah Field and sought to examine whether stockpile age impacted vegetation establishment in a greenhouse setting. We hypothesized that vegetation establishment would be different between stockpile ages. Additionally, we hypothesized soil stockpile age would impact seed head formation on vegetation within our study.

## Methods

### Study Area

All soils in this study were collected from the Jonah Infill natural gas field in Sublette County, WY, USA (Figure 1). The Jonah Infill natural gas field is located at 2,100m, has an average of 39-52 frost free days per year, and receives an average of 17.8-22.9 cm (7-9 inches) precipitation per year. A total of 6514 acres have been disturbed in the Jonah Field in the form of well pads, pipelines, and roadways with a total of 4817 acres (74%) being under reclamation. Well pads in the Jonah Field average ∼5.3 acres in size.

**Figure 1.**
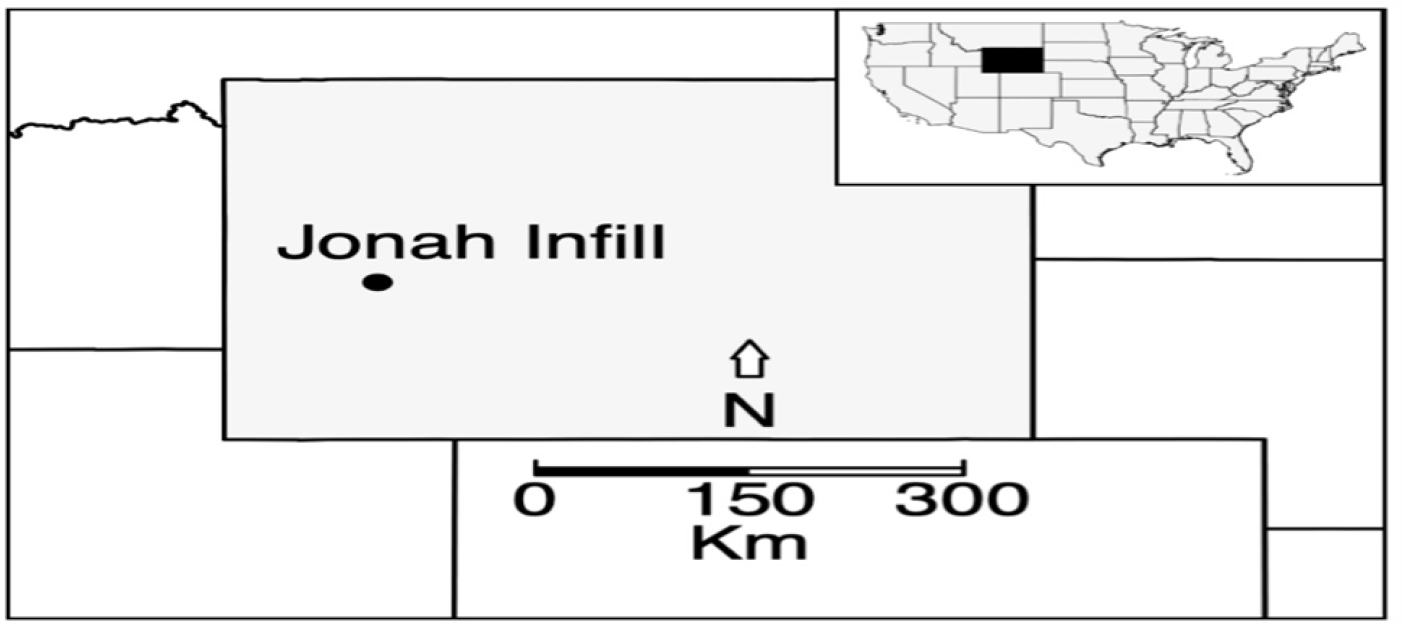
A map depicting the location of the Jonah Infill natural gas field within the state of Wyoming.

### Soil Handling and Collection

All natural gas well pads in this study are in Ecological Site Description R034AC126WY which consists of loamy soils. In the Jonah Field, suitable topsoil is stripped and stockpiled adjacent to the well pad prior to well pad construction. Stockpiles remain in place until natural gas exploration activities are complete, at which time stockpiled soil is respread to initiate interim reclamation. Stockpiles in the Jonah Field are seeded with the same seed mix as used for reclamation in effort to prevent erosion, combat weeds and to keep soil processes active.

In this study, soils from 17 total stockpiles were utilized with three replicates each from stockpiles aged 1-5 years old, and one each of 6- and 7-year-old stockpiles. Soil from each stockpile was removed from roughly 1.5m (5 feet) below the surface of the stockpile using a backhoe and subsequently placed and sealed into 5-gallon buckets in the Jonah Field on 30 October 2019. After soil was collected, it was transported in sealed 5-gallon buckets to Laramie, WY where it was placed in a freezer at a University of Wyoming Agriculture Experimental Station for later testing to best simulate the Jonah Field which is typically freezing throughout winter months. Two weeks prior to seeding, soil was removed from the freezer and placed outside to thaw out.

### Study Design and Seeding

This study utilized a split-plot design where age class was the whole-plot treatment factor (3 replicates of the 1–5-year-old age classes and a single replicate for the 6- and 7-year age class) and the subplot factor as time. Each replicate was sub-divided into four pseudo-replicates (12.7 cm (5 inch) diameter plastic pots) and the pots were randomly placed within a 68-block grid (6 rows of 10 and 1 row of 8, Figure 2) within a greenhouse controlled at 21.1°C (70°F) in the daytime and 18.3°C (65°F) at night. Each pot was watered during the initial placement into the greenhouse and was not watered further in the study. Each pot was seeded with a native seed mix commonly used in the Jonah Field on June 23, 2020. The seed mix contained 6 grasses which are commonly drill seeded, as well as 9 forbs and 4 shrubs which are typically broadcast seeded and raked over (see Appendix A for seed mix information). To mimic this, after the pots were filled 3/4 of the way with soil, grass seed was spread over the soil and an additional 1/4 inch of soil was added. Then, forbs and shrubs were added and lightly raked over with a plastic fork.

**Figure 2.**
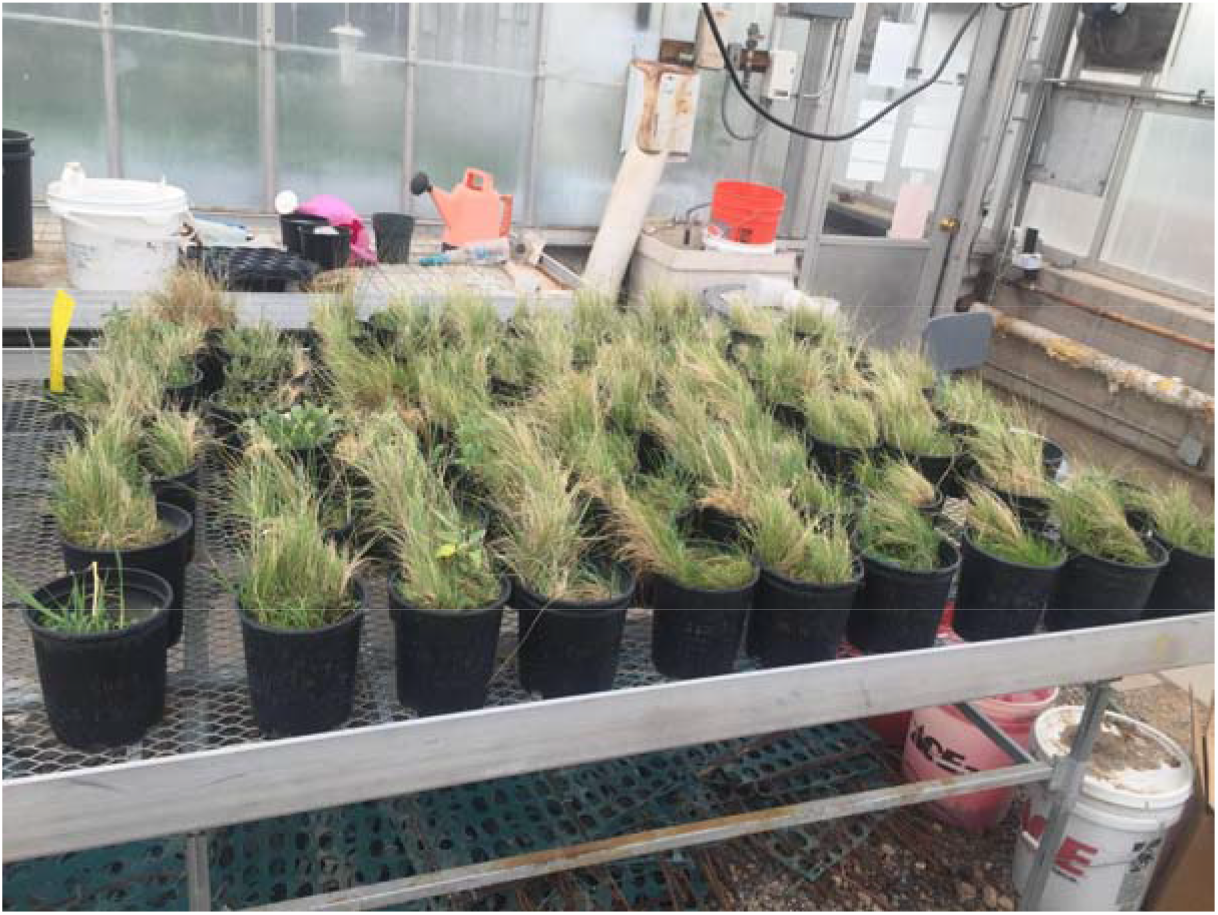
An image showing the layout of the study design within a greenhouse at the University of Wyoming Agricultural Experiment Station.

### Vegetation Sampling

Digital images were taken above the vegetation starting four weeks after the seeding date and subsequently every 2 weeks thereafter, totaling four sets of images, with the last set being taken slightly 10 weeks after initial seeding. Nadir images were taken at 0.7 meters above each pot with a 14-megapixel Kodak Easyshare M532 handheld camera (Eastman Kodak Corporation, Rochester, NY, USA). This resulted in a ground sample distance (GSD) of ∼0.25 mm which is similar to imagery in previous peer-reviewed studies to assess vegetation on reclaimed sites (e.g., Curran et al. 2019, 2020). All images were cropped to isolate individual pots and four photos representing each stockpile (one per each pot) were placed into folders labeled by the stockpile name and date of photo. Photos were then analyzed with a free image analysis software called ‘SamplePoint’ with 25 pixels being analyzed per image using a 5×5 grid (Booth et al. 2006). This resulted in 25 pixels being sampled within each pot, or 100 pixels over the four pots per each stockpile during each photo period. Pixels within each image were classified by setting ‘buttons’ in SamplePoint to ‘vegetation’ or ‘soil’ across each image set. The total values were converted into percent cover for each category (e.g., if 24 of 25 pixels were classified as ‘vegetation’ and 1 of 25 pixels was soil, the pot was considered to have 96% vegetation cover and 4% soil cover). The images taken on the last day of the study were also assessed for seed head appearance.

### Statistical Analysis

To assess differences in vegetation establishment, the median percent vegetation cover across the four plastic pots for each stockpile/age class combination was measured 4 weeks, 6 weeks, 8 weeks, and 10 weeks after the initial planting. For age classes 1 year – 5 years, there were a total of 60 observations (3 stockpiles * 5 years * 4 time periods) and for age classes 6 and 7 years, there were a total of 8 observations (1 stockpile * 2 years * 4 time points). Statistical analyses were conducted using the nlme library of the R statistical programming language (R Core Team 2020). A repeated measures analysis of variance was used to model percent vegetation cover as a function of age class, time, and the interaction of age class and time.

## Results

There was a statistically significant increase in percent vegetation cover across time (p<0.0001 with a mean percent increase of 10.8% in cover every 2 weeks (Figure 3). However, the mean increase over time was not statistically different among the age-classes (p=0.1408, Figure 3). Percent cover did not differ significantly among the age classes at any measuring event (p>0.26 for every pairwise comparison of age classes). Additionally, while statistical analyses were not conducted on seed head presence, all 68 pots within the study contained vegetation with seed heads present at the 10-week sampling period.

**Figure 3.**
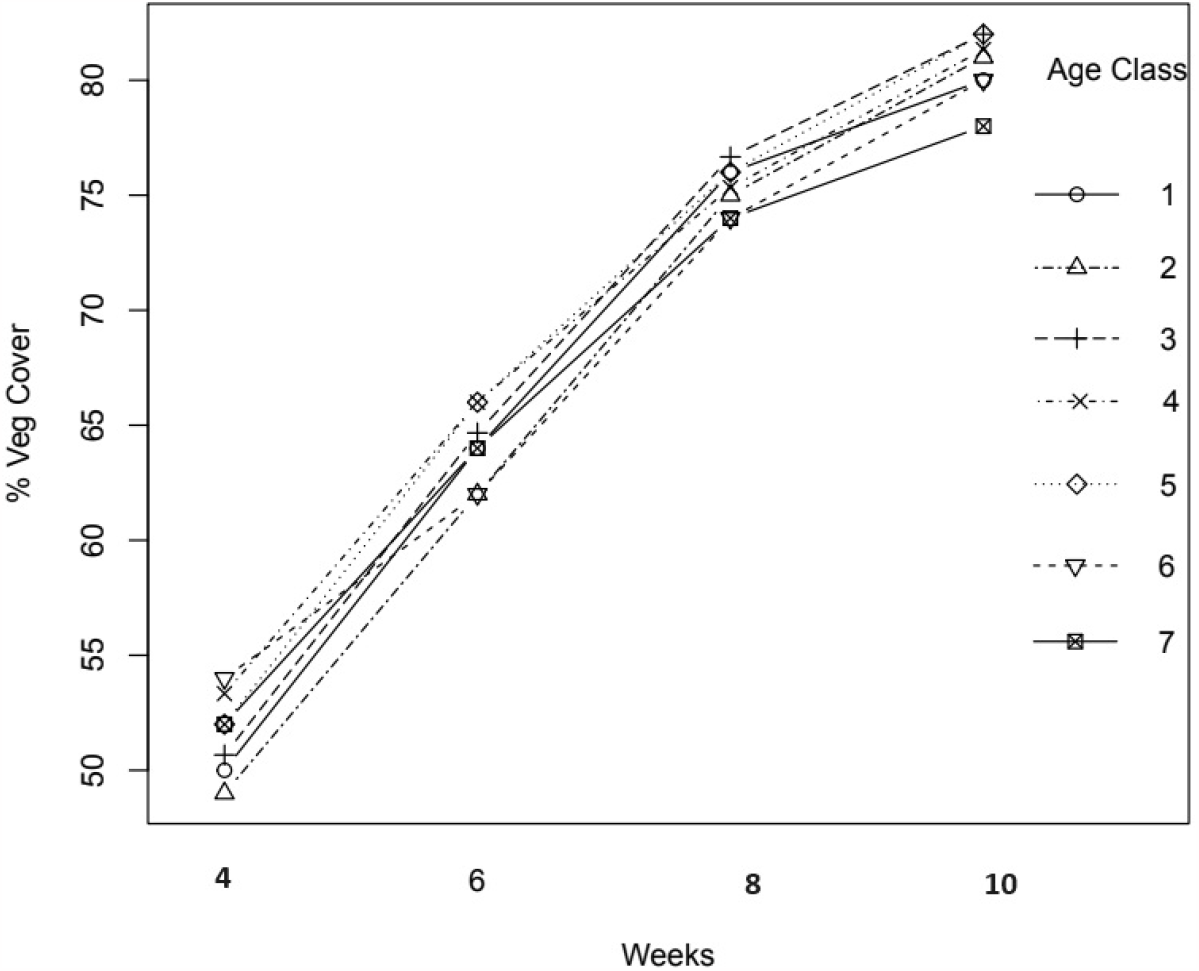
A graphical representation of median percent vegetation cover per age class across the duration of this study. Vegetation cover increased over time but was not significantly different among age classes at any sampling period.

## Conclusion

Our results do not support our hypotheses that soil stockpile age impacts vegetation establishment or ability to set seed. Rather, they indicate that vegetation establishment and ability to produce seed are not determined by stockpile age. Vegetation establishment is critical in ensuring a critical short-term reclamation goal of preventing erosion. Seed production is critical in ensuring a self-sustaining vegetation community to achieve longer-term goals (Herrick et al. 2006). Therefore, our results are likely to have important management implications. Specifically, the addition of amendments, fertilizers or other biotic stabilizers may not be necessary to reestablish native vegetation within the Jonah Field following development. Rather, planting native vegetation on soil stockpiles is sufficient to maintain soil health.

Previous research has suggested most soil degradation associated with stockpiles in oil and gas fields is a result of initial surface disturbance and respreading soil rather than when soil is maintained in the stockpile (Mason et al. 2011). As stockpiles in the Jonah field are seeded after being created, it is likely carbon inputs from plants help maintain soil biotic functionality (Valliere et al. 2022). Previous research has shown additional disturbance to soil after stockpiling and respreading exacerbates soil degradation (Wick et al. 2009, Mason et al. 2011). With the advent of directional drilling, it may not always be certain when drilling will be complete on a given well pad since additional wells may be placed on a given well pad if flowback rates are high during exploratory drilling. Therefore, if vegetation establishment is unaffected by soil stockpile age, it is likely more beneficial to keep soil in stockpiles until disturbances associated with drilling are complete than it is to respread soil with potential for redisturbance. Not only does redisturbance exacerbate soil degradation, but it may also lead to destruction of successful reclamation.

Like other greenhouse studies, our study has limitations of being short-term and in a controlled environment. However, our results clearly show soil from the Jonah Infill natural gas field remains fertile in stockpiles for at least 7 years. This study would be enhanced by additional, longer-term studies in field conditions. Furthermore, since stockpiles greater than 7 years old were not available, additional research to determine at what age, if any, soil fertility is lost in stockpiles in this environment. Other areas of stockpile research which may benefit operators and reclamationists associated with oil and gas development would be to expand a study like this across various climate regimes, as it is possible stockpiles act as cold storage areas in the Jonah Infill due to its extreme climatic conditions. While the purpose of this study was to examine vegetation emergence in different aged stockpiles, there may be benefit in examining soil physical and chemical properties in different aged soil stockpiles over time. Finally, while many species were seeded in this study, this study only analyzed percent vegetation cover because species specific information was difficult to obtain until seed heads emerged late in the study and because it was not possible to have an exactly equal amount of seed from each species in each pot as seed was obtained in pre-mixed bags. Future studies to determine if specific vegetation species are impacted by soil stockpile age may be useful.

## About the Authors

Michael Curran owns and operates Abnova Ecological Solutions. He received his PhD in Ecology from the University of Wyoming in 2020. He received the M.S. Student Scholarship from the American Society of Mining and Reclamation in 2014 and the PhD Student Scholarship from the American Society of Reclamation Sciences in 2020. Mike is a Certified Ecological Restoration Practitioner and a Certified Wildlife Biologist and serves as Editor of *Reclamation Matters* and Associate Editor for *Reclamation Sciences* and *Natural Areas Journal*.

Joshua Sorenson is the Senior Reclamation Specialist for Jonah Energy in Pinedale, WY and holds an M.S. degree from Texas A&M University. He received the 2021 Reclamationist of the Year award from the American Society of Reclamation Sciences. Josh manages reclamation on over 1,900 well pads for Jonah Energy and has helped advance the science and practice of reclamation in the State of Wyoming, as he has successfully reclaimed hundreds of acres of lands in an environment which has less than a 50-day growing season and receives under 10 inches of annual precipitation.

Tim Robinson is a Professor of Statistics at the University of Wyoming. He is a Fellow of the American Statistical Association and the American Society for Quality. His areas of research include applications of statistics to ecology, the environment, engineering, and medicine.

## Appendix A

Seed mix used for reclamation in the Jonah Infill natural gas field.

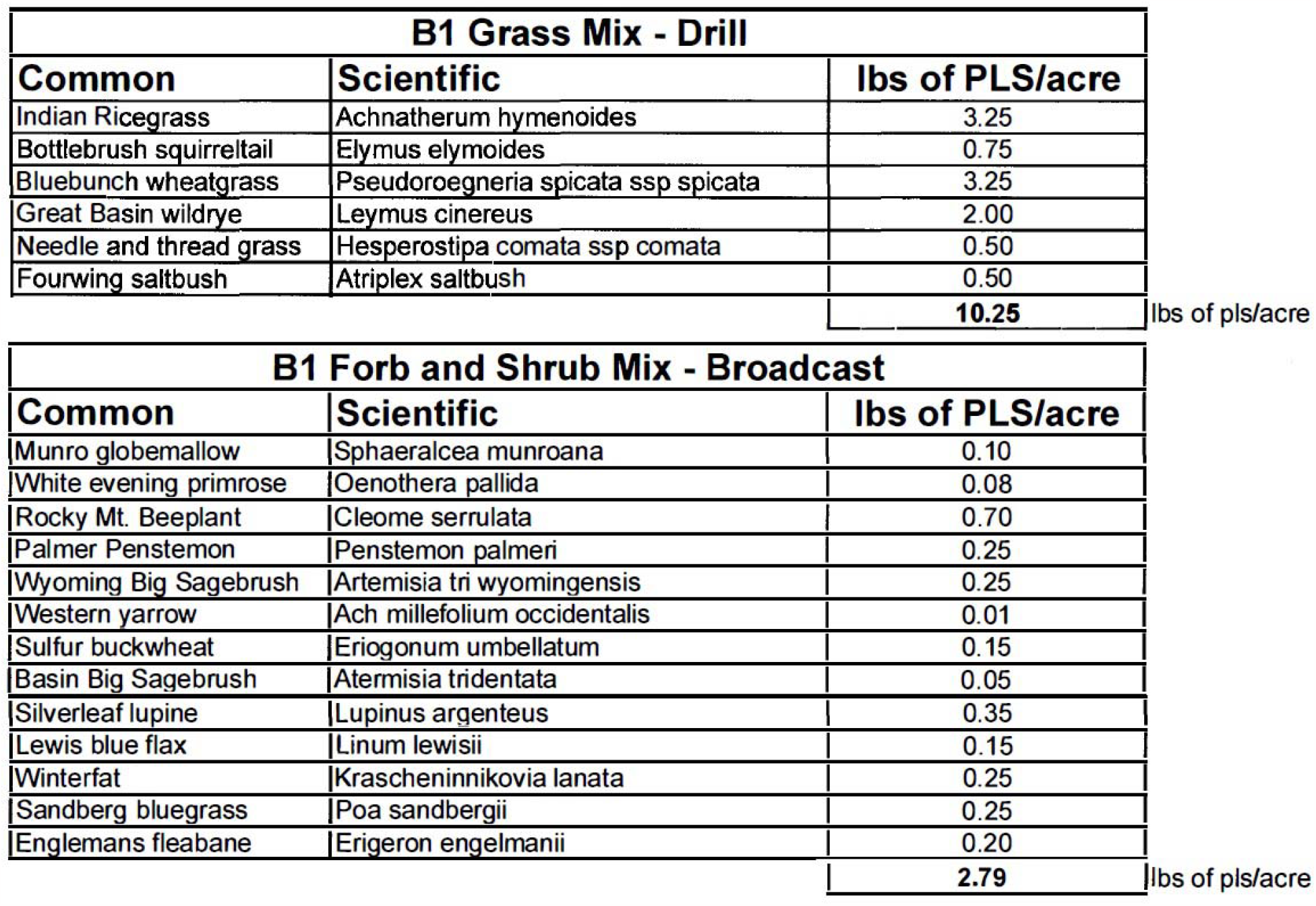

