## Appendix A for "Soil stockpile age does not impact vegetation establishment in a cold, arid natural gas field"

**Supplemental Material**

**Appendix A**

Seed mix used for reclamation in the Jonah Infill natural gas field.

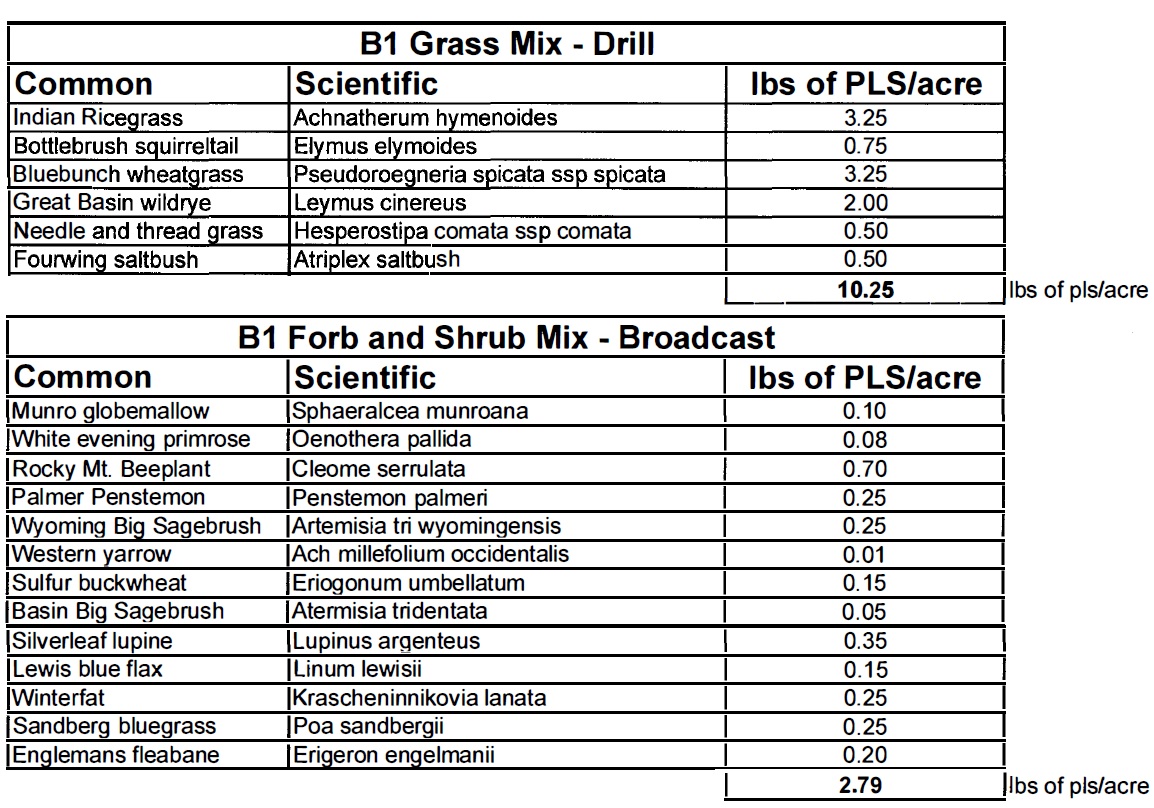
